# A transcriptome for the early-branching fern *Botrychium lunaria* enables fine-grained resolution of population structure

**DOI:** 10.1101/2020.02.17.952283

**Authors:** Vinciane Mossion, Benjamin Dauphin, Jason Grant, Niklaus Zemp, Daniel Croll

## Abstract

Ferns are the second most dominant group of land plants after angiosperms. Extant species occupy an extensive range of habitats and contribute significantly to ecosystem functioning. Despite the importance of ferns, most taxa are poorly covered by genomic resources. The genus *Botrychium* belongs to the family Ophioglossaceae, one of the earliest divergent lineages of vascular plants, and has a cosmopolitan distribution with 35 species, half of which are polyploids. Here, we establish a transcriptome for *Botrychium lunaria*, a diploid species with an extremely large genome with a 1C value of 12.10 pg. We assembled 25,701 high-quality transcripts with an average length of 1,332 bp based on deep RNA-sequencing of a single individual. We sequenced an additional 11 transcriptomes of individuals from two populations in Switzerland, including the population of the reference individual. Based on read mapping to reference transcript sequences, we identified 374,510 single nucleotide polymorphisms (SNPs) segregating among individuals for an average density of 14 SNPs per kb. The transcriptome-wide markers provided unprecedented resolution of the population genetic structure revealing substantial variation in heterozygosity among individuals. We also constructed a phylogenomic tree of 90 taxa representing all fern orders to ascertain the placement of the genus *Botrychium*. The high-quality transcriptomic resources enable powerful population and phylogenomic studies in an important group of ferns.

**Significance statement:** Ferns pose substantial puzzles in terms of lifestyles, genome organization and population structure. Progress has been significantly hampered by the lack of genomic resources. Here, we present a transcriptome for *Botrychium lunaria*, a phylogenetically early-branching fern with an extremely large genome. We show that the new transcriptome improves phylogenetic resolution among early-branching ferns. Based on an additional 11 transcriptomes of the same species, we identify unexpected variation in population-level heterozygosity.

## Introduction

Ferns (Polypodiopsida) constitute the earliest divergent lineage of vascular plants along with lycophytes (Lycopodiopsida) (Kranz and Huss, 1996; Kenrick and Crane, 1997). With 85% of the total species richness found in the tropics, ferns are present in most climates (Page, 2002; Ranker and Haufler, 2008). Habitats of ferns include deserts, grasslands, forest understory, mountainous regions, and aquatic environments (Mehltreter *et al*., 2010) where they have diversified into a multitude of lifestyles. Ferns play key roles in ecosystem functioning including serving as a habitat for invertebrates (Ellwood and Foster, 2004), shaping plant recolonization of disturbed habitats (Walker, 1994) and influencing the composition of tree species communities (George and Bazzaz, 1999a; George and Bazzaz, 1999b). Ferns have complex and idiosyncratic life cycles. In contrast to other plants, many fern species are capable of versatile reproductive modes (Sessa *et al*., 2012) including apomixes, sporophytic and gametophytic selfing, as well as outcrossing (Barker and Wolf, 2010; Haufler *et al*., 2016). Yet, our understanding of the evolutionary origins and lifestyle diversification is limited.

Phylogenetic analyses resolved the position of ferns as the sister group to seed plants (Pryer *et al*., 2001) and as the second earliest diverging lineage of vascular land plants (Raubeson and Jansen, 1992; Pryer *et al*., 2004). The crown age of ferns was estimated to be ca. 360-431 million years underlining the deep divergence among ferns lineages (Pryer *et al*., 2004; Testo and Sundue, 2016; Rothfels *et al*., 2015; Lehtonen *et al*., 2017; Magallón *et al*., 2013; Des Marais *et al*., 2003; Zhong *et al*., 2014; Wikström and Kenrick, 2001; Qi *et al*., 2018). Fern phylogenies have been established based on chloroplast markers (Schuettpelz and Pryer, 2007; Rai and Graham, 2010; Kuo *et al*., 2011; Lu *et al*., 2015; Testo and Sundue, 2016; Grewe *et al*., 2013; Lu *et al*., 2015; Rai and Graham, 2010) in combination with mitochondrial (Knie *et al*., 2015) and nuclear markers (Pryer *et al*., 2001; Pryer *et al*., 2004; Schuettpelz *et al*., 2016) or both (Qiu *et al*., 2007). Plastid and mitochondrial markers are unsuitable to investigate events of reticulate evolution, whereas a third of all speciation events may have been driven by polyploidization (Wood *et al*., 2009). In contrast, a series of recent phylogenomic studies highlighted the power of transcriptome-based approaches (Wickett *et al*., 2014; Shen *et al*., 2018; Leebens-Mack *et al*., 2019; Qi *et al*., 2018; Rothfels *et al*., 2015; Rothfels *et al*., 2013). Expanding genome- or transcriptome-wide datasets will help to further improve the accuracy of phylogenomic reconstructions.

The first insights into the structure of fern genomes were provided by two complete genome sequences of *Azolla filiculoides* and *Salvinia cucullata* (Li *et al*., 2018). These two species belong to the 1% heterosporous fern species in contrast to the dominant homosporous species. Heterosporous ferns exhibit the smallest known genome sizes as shown for the closely related species *A. microphylla* and *S. molesta* (1C value of 0.77 pg and 1C value of 2.28 pg, respectively; Obermayer *et al*., 2002; Clark *et al*., 2016). Conversely, homosporous fern genomes contain on average three times more chromosomes than heterosporous ferns and seeds plants (Barker and Wolf, 2010). The recent release of a partial genome assembly for the heterosporous fern *Ceratopetris richardii* highlighted the challenge associated with complex fern genomes (Marchant *et al*., 2019). Major progress in the establishment of genomic resources was made with the sequencing of 73 fern transcriptomes (Leebens-Mack *et al*., 2019; Carpenter *et al*., 2019). Such datasets were successfully used to develop single copy nuclear markers to resolve deep evolutionary relationships among ferns (Rothfels *et al*., 2013; Rothfels *et al*., 2015). Transcriptome assemblies are also an important tool to develop genotyping approaches and overcome challenges associated with extremely large fern genome (Bennett and Leitch, 2001; Obermayer *et al*., 2002; Hanson and Leitch, 2002). These approaches typically reduce genome complexity but still provide sufficient polymorphic markers to conduct population genomics analyses (Seeb *et al*., 2011). Establishing transcriptomic datasets for understudied fern clades will bring new insights into fern diversification.

An important genus lacking transcriptomic resources is *Botrychium* belonging to subclass Ophioglossidae (PPG I, 2016). This subclass is characterized by a subterranean gametophytic stage (Jeffrey, 1898; Winther and Friedman, 2007; Field *et al*., 2015) and extremely large and complex genomes (e.g., *Ophioglossum petiolatum* 1C value of 65.55 pg; Obermayer *et al*., 2002). *Botrychium* occurs in open habitats on nearly every continent across a broad temperate and boreal distribution. This genus is divided into three monophyletic clades defined by maternally inherited markers (Simplex-Campestre, Lanceolatum and Lunaria; Dauphin *et al*., 2017), containing 35 recognized taxa (PPG I, 2016). The challenge of identifying *Botrychium* taxa based on morphology is underlined by claims of cryptic species (Clausen, 1938; Hauk, 1995). Ambiguous morphologies are sometimes caused by polyploidization which is a major driver of speciation as half of the known *Botrychium* species are allopolyploids (Dauphin *et al*., 2018). Nuclear markers resolved the parental origins of these allopolyploid taxa and provided insights into the genus radiation approximately 2 million years ago. Additionally, the reconstruction of maternal lineages of *Botrychium* revealed genetic diversity within the Lunaria clade highlighting the uncertainty of taxonomic assignments (Dauphin *et al*., 2014; Maccagni *et al*., 2017; Dauphin *et al*., 2017). Previous population genetic studies based on isozymes showed a lack of genetic differentiation among morphologically recognized types (Williams *et al*., 2016), and the low amount of genetic variation detected within *Botrychium* populations suggests pervasive self-fertilization (Farrar, 1998; Hauk and Haufler, 1999). Furthermore, genetic differentiation among populations and regions was found to be low suggesting that gene flow may occur (Camacho and Liston, 2001; Swartz and Brunsfeld, 2002; Birkeland *et al*., 2017). These studies highlight the need for powerful, genome-wide marker systems to resolve population structures, life histories and taxonomy of these early-branching ferns.

In this study, we assemble and curate the transcriptome of *B. lunaria* with a massive genome size of a 1C value of 12.10 pg (Veselý *et al*., 2012) filling an important gap in the fern phylogeny. By analyzing the transcriptome of an additional 11 individuals, we show that individuals vary substantially in terms of genotype and heterozygosity within and between populations. We further demonstrate the power of transcriptome-wide markers to resolve phylogenetic relationships at the genus level and among deeply divergent fern lineages.

## Material and Methods

### Sampling, library preparation and sequencing

Leaf material of *B. lunaria* was obtained from three locations in Switzerland: two in the Valais Alps in Val d’Hérens, Mase and Forclaz within approximately 30 km, and one in the Jura Mountains at Chasseral (Table 1). Leaves of six individuals from Val d’Hérens and from Chasseral were collected in July 2015 and June 2017, respectively. Plant material was wrapped in aluminum foil and frozen immediately in liquid nitrogen. Total RNA was extracted from trophophores (i.e., sterile part of leaves) using the RNAeasy Plant Mini Kit (Qiagen) and DNA was eliminated using DNase I digestion. Total RNA was quantified using a Qubit fluorometer (Invitrogen, Thermo Fisher Scientific) with the RNA Broad-Range assay kit (Invitrogen, Thermo Fisher Scientific) and quality-checked using an Agilent 2200 Tape Station (Agilent Technologies, Inc.). Samples were diluted to 100 ng/μl in RNase free ultra-pure water before library preparation. The RNA-sequencing libraries were prepared following a TruSeq RNA library preparation protocol (Illumina, Inc.) enriching for polyadenylated RNAs. After quality assessment on an Agilent 2200 Tape Station, libraries were pooled and sequenced in 150 bp single-end mode on one lane of an Illumina HiSeq 4000 sequencer.

**Table 1:** Populations, accessions and voucher information.

| Individual identifier | Population | Location | Latitude | Longitude | Altitude (m) | Date | Voucher number | Deposit<sup>1,2</sup> institute |
| --- | --- | --- | --- | --- | --- | --- | --- | --- |
| CHA_I_1 | Chasseral | Chasseral | 47.12974 | 7.04934 | 1549.71 | 07.06.2017 | NE000101258 | NE |
| CHA_I_3 | Chasseral | Chasseral | 47.12974 | 7.04934 | 1549.71 | 07.06.2017 | NE000101257 | NE |
| CHA_I_5 | Chasseral | Chasseral | 47.12974 | 7.04934 | 1549.71 | 07.06.2017 | NE000101256 | NE |
| CHA_I_6 | Chasseral | Chasseral | 47.12974 | 7.04934 | 1549.71 | 07.06.2017 | NE000101255 | NE |
| CHA_I_7 | Chasseral | Chasseral | 47.12974 | 7.04934 | 1549.71 | 07.06.2017 | CHA_I_7 | UNINE |
| CHA_I_8 | Chasseral | Chasseral | 47.12974 | 7.04934 | 1549.71 | 07.06.2017 | NE000101254 | NE |
| IIT2_H2 | Val d'Hérens | Forclaz | 46.08741 | 7.54379 | 2346.15 | 2015 | IIT2_H2 | UNINE |
| IIT1_H5 | Val d'Hérens | Forclaz | 46.08851 | 7.53906 | 2346.15 | 2015 | IIT1_H5 | UNINE |
| IIT2_B2 | Val d'Hérens | Forclaz | 46.08741 | 7.54379 | 2406.10 | 2015 | IIT2_B2 | UNINE |
| IT1_A3 | Val d'Hérens | Mase | 46.20432 | 7.48350 | 2406.62 | 2015 | IT1_A3 | UNINE |
| IT1_H1 | Val d'Hérens | Mase | 46.20432 | 7.48350 | 2406.62 | 2015 | IT1_H1 | UNINE |
| IT2_D4 | Val d'Hérens | Mase | 46.19642 | 7.48502 | 2424.07 | 2015 | IT2_D4 | UNINE |

Notes: <sup>1</sup>NE: Herbarium of the University of Neuchâtel, <sup>2</sup>UNINE: University of Neuchâtel, Evolutionary genetics laboratory.
The specimens deposited at UNINE are frozen samples stored at -80°C.

### De novo assembly, filtering and quality assessment

Sequencing reads were quality-checked using FastQC v. 0.11.7 (Andrews, 2010) and trimmed using Trimmomatic v. 0.38 (Bolger *et al*., 2014). Reads were retained if the leading and trailing bases > 5, a 4-bp sliding window > 15, and a minimum read length of 36 bp. Trimmed transcript sequences were then *de novo* assembled using Trinity v. 2.8.3 (Haas *et al*., 2013) from a single individual used as a reference. We used the pseudo-alignment percentage calculated by Kallisto v. 0.45.0 (Bray *et al*., 2016) to assess the representativeness of the raw assembly across the twelve sequenced individuals in total. Candidate coding regions were identified using TransDecoder v. 5.3.0 (Haas *et al*., 2013). Only transcripts with an open reading frame (ORF) of at least 100 amino acids were kept. We also retained only the longest isoform per transcript using the Trinity v. 2.8.3 toolkit. We used Diamond v. 0.9.24 (Buchfink *et al*., 2015) to screen the transcript assembly against the NCBI non-redundant protein (nr) and UniVec databases to identify potential foreign RNA contaminants. The best hit for each transcript was assigned at the phylum-level using the R package *taxise* v. 0.9.7 (Chamberlain and Szöcs, 2013) in RStudio v. 1.2.1335 (RStudio Team, 2015; R Development Core Team, 2020). We excluded all transcripts with a best hit outside of the plant kingdom. The completeness of the transcriptome assembly was assessed using BUSCO v. 4.0.6 with the viridiplantae_odb10 database (Simão *et al*., 2015). Data were visualized using the R package *ggplot2* v. 3.2.1 (Wickham, 2016).

### Variant calling

We generated alignment BAM files for each individual against the transcriptome using the short read aligner Bowtie2 v. 2.3.5 (Langmead, 2010) and SAMtools v. 1.9 (Li *et al*., 2009). Depth coverage of the reference individual was extracted using SAMtools idxstats. Alignments were processed with HaplotypeCaller implemented in the Genome Analysis Toolkit (GATK) v. 4.1.0.0 (DePristo *et al*., 2011; McKenna *et al*., 2010; Van der Auwera *et al*., 2013) for single nucleotide polymorphism (SNP) calling. The resulting gvcf files were combined and genotyped using the GATK CombineGVCF and GenotypeGVCF tools, respectively. We excluded monomorphic sites from further analysis. We filtered SNPs for the number of genotyped chromosomes (AN ≥ 20) out of a maximum of 24 (12 diploid individuals). Quality criteria QUAL > 100, QualByDepth > 5.0, RMSMappingQuality > 20.0, MappingQualityRankSumTest retained values > −2.0 and < 2.0, and ReadPosRankSumTest and BaseQualityRankSumTest retained values > −2.0 and < 2.0 were defined following the best practices and were applied to flag low-quality loci (Suppl. Figure S1). We removed SNPs failing the above filters using VCFtools v. 0.1.16 (Danecek *et al*., 2011) and added a filter to retain only bi-allelic SNPs. These analyses were performed using the R packages *vcfR* v. 1.10.0 (Knaus and Grünwald, 2017) and the SNP statistics among transcripts were visualized using *ggplot2* v. 3.2.1.

### Population genetics analyses

Intra-individual allele frequencies were calculated for each individual and SNP locus using the mapped read depth per allele (AD). The frequency distributions were plotted per individual. Then, to avoid biases introduced by highly polymorphic transcripts and to reduce the computational time of population genetic analysis, we subsampled the number of SNPs by selecting one SNP every 1,000 bp of transcriptomic sequence using VCFtools v. 0.1.16. We performed principal component analyses (PCA) and calculated the pairwise *F*ST and the mean heterozygosity (*H*e) per location and per individual. These analyses were performed using the R packages *vcfR* v. 1.10.0, *adegenet* v. 2.1.3 (Jombart and Ahmed, 2011) and *hierfstat* v. 0.04-22 (Goudet, 2005), and data were visualized using *ggplot2* v. 3.2.1.

### Functional annotation

We functionally characterized encoded protein sequences based on gene ontology (GO) terms. We summarized GO terms by selecting the least redundant annotations among the 30 most frequent terms per ontology (cellular component CC, molecular function MF, and biological process BP). Analyses were performed using the Bioconductor packages *AnnotationDbi* v. 1.46.0 (Pagès *et al*., 2020), *GO*.*db* v. 3.8.2 (Carlson, 2020), *GSEABase* v. 1.34.0 (Morgan *et al*., 2020), *annotate* v. 1.62.0 (Gentleman, 2020), and data were visualized using the R packages *ggplot2* v. 3.1.0.

### Genus-level phylogenetic analyses

We analysed published sequences of four nuclear regions from diploid and polyploid *Botrychium* taxa: *ApPEFP_C, CRY2cA, CRY2cB*, and *transducin* (Dauphin *et al*., 2018). We searched homologous sequences in the transcriptome assembly using BLAST v. 2.9.0 (Altschul *et al*., 1990). If the associated transcript was found in the assembly, we used BCFtools v. 1.9 (Li, 2011) to retrieve the corresponding transcript from the 11 remaining individuals using the VCF file information. Sequence alignments were performed with MAFFT v. 7.470 under the G-INS-i strategy and default parameters (Katoh *et al*., 2002; Katoh and Standley, 2013). Multiple alignments were visually inspected and manually adjusted using Geneious v. 8.1.9 (Kearse *et al*., 2012). Phylogenetic trees were inferred using maximum likelihood (ML) in RAxML-NG v. 0.9.0 (Kozlov *et al*., 2019). We ran tree inferences with a fixed random seed of 2 under the HKY+GAMMA model based on model settings by Dauphin *et al*. (2018) to ensure reproducibility. The tree search was performed using 25 random- and 25 parsimony-based starting trees. The branch support was estimated using 1,000 bootstrap replicates and calculated according to the transfer bootstrap expectation matrix (Lemoine *et al*., 2018). The support values were depicted on the best-scoring ML tree. Final gene trees were edited with FigTree v. 1.4.4 (Rambaut, 2009).

### Phylogenomic analyses

We performed phylogenomic analyses across ferns by including the newly established *B. lunaria* transcriptome for a total of 95 transcriptomes including 86 fern species (18 Eusporangiates, 68 Leptosporangiates), six Spermatophyta and two Lycopodiopsida (Suppl. Table S1; Shen *et al*., 2018; Qi *et al*., 2018; Leebens-Mack *et al*., 2019). The Spermatophyta and Lycopodiopsida species represent outgroups in the analysis. We first performed an ortholog search using OrthoFinder v. 2.3.12 (Emms and Kelly, 2015) including the newly established *B. lunaria* transcriptome and protein sequences of other transcriptomes. We retained only orthogroups with members found in at least 85 species (> 90% of total), with a minimum of 80 species (> 85%) carrying a single copy ortholog. Species represented in less than 50% of the orthogroups were excluded from the dataset. Members of each orthogroup passing our filters had multiple gene copies per individual. Then, we randomly selected one copy of each duplicated gene (i.e., the first copy reported by OrthoFinder) to reduce the amount of missing data in gene trees. Sequences of the orthogroups subset were subsequently aligned with MAFFT v. 7.470 under the L-INS-i strategy and default parameters. The optimal substitution model was assessed for each orthogroup alignment using modeltest-ng v. 0.1.6 (Darriba *et al*., 2020). Finally, unrooted gene trees were built using maximum likelihood (ML) in RAxML-NG v. 0.9.0. We ran tree inferences under the best model according to the Akaike information criterion (AICc) criterion with a fixed random seed of 2. The tree search was performed on 25 random and 25 parsimony-based starting trees and branch support was estimated over 100 bootstrap replicates. The inferred gene trees were used to estimate a species tree with Astral v. 5.7.3 (Mirarab *et al*., 2014; Zhang *et al*., 2018). Branch support was calculated using multi-locus bootstrapping (Seo, 2008) and local posterior probabilities (Sayyari and Mirarab, 2016). Species trees were edited with FigTree v. 1.4.4 (Rambaut, 2009). Phylogenetic trees, alignments and protein sequences are available as Supplementary Files S1-S5 https://doi.org/10.5281/zenodo.3959727.

## Results

### Sample collection and transcriptome assembly

In total, twelve *B. lunaria* individuals were sampled from Val d’Hérens and Chasseral located in the Valais Alps and the Jura Mountains, respectively. The two sites are approximately 120 km apart and the Alpine population was sampled on meadows on an altitudinal range of 1,500 to 2,400 meters (Table 1). The transcriptome sequencing produced 14.6-50.1 million reads per individual. After quality trimming, we retained 97.0-99.2% of the reads (Figure 1A, Suppl. Table S2). The highest number of high-quality reads (49.5 million) were obtained for the Chasseral individual CHA_I_1. Hence, we selected this individual to produce a transcriptome. The raw assembly for the reference individual (CHA_I_1) contained 167,306 transcripts for a total of 87,537 candidate genes. Mapping reads from individuals to the raw transcriptome assembly showed an alignment rate (percentage of mapped reads) between 74.2-82.5% regardless of the population of origin (Figure 1A, Suppl. Table S2). We analyzed all transcripts for the presence of high-confidence open reading frames (ORF; ≥100 amino acids). We retained 69,280 transcripts (41.4%) covering 26,139 predicted genes (Figure 1B). Next, we selected the longest transcript for each gene (Figure 1D). We performed a screen of each gene against the complete non-redundant protein and the UniVec database of NCBI and found evidence for contamination in 438 transcripts. Most contaminant sequences were associated with viruses, fungi and bacteria (Figure 1C). The final assembly consisted of 25,701 unique transcripts spanning a total of 34.24 Mb. The average and median transcript length were 1,332 and 967 bp, respectively (Figure 1D). The N50 of the final transcriptome was 1,995 bp with an average GC content of 44.3% (Table 2). GO terms were assigned to 12,154 transcripts (47.2%; Figure 2).

**Table 2:** Overview of assembly statistics over the different transcript filtering stages.

| Filtering stage | Assembled bases | Transcripts | Genes | N50-longest isoform | N50-all | GC% |
| --- | --- | --- | --- | --- | --- | --- |
| Raw assembly | 56,273,802 | 167,306 | 87,537 | 1,689 | 1,089 | 43.68 |
| ORF-encoding | 34,588,465 | 69,280 | 26,139 | 2,152 | 1,988 | 44.00 |
| Longest isoform only | 34,588,465 | 26,139 | 26,139 | 1,988 | 1,988 | 44.31 |
| Contaminants screening | 34,245,455 | 25,701 | 25,701 | 1,995 | 1,995 | 44.30 |

**Figure 1:**
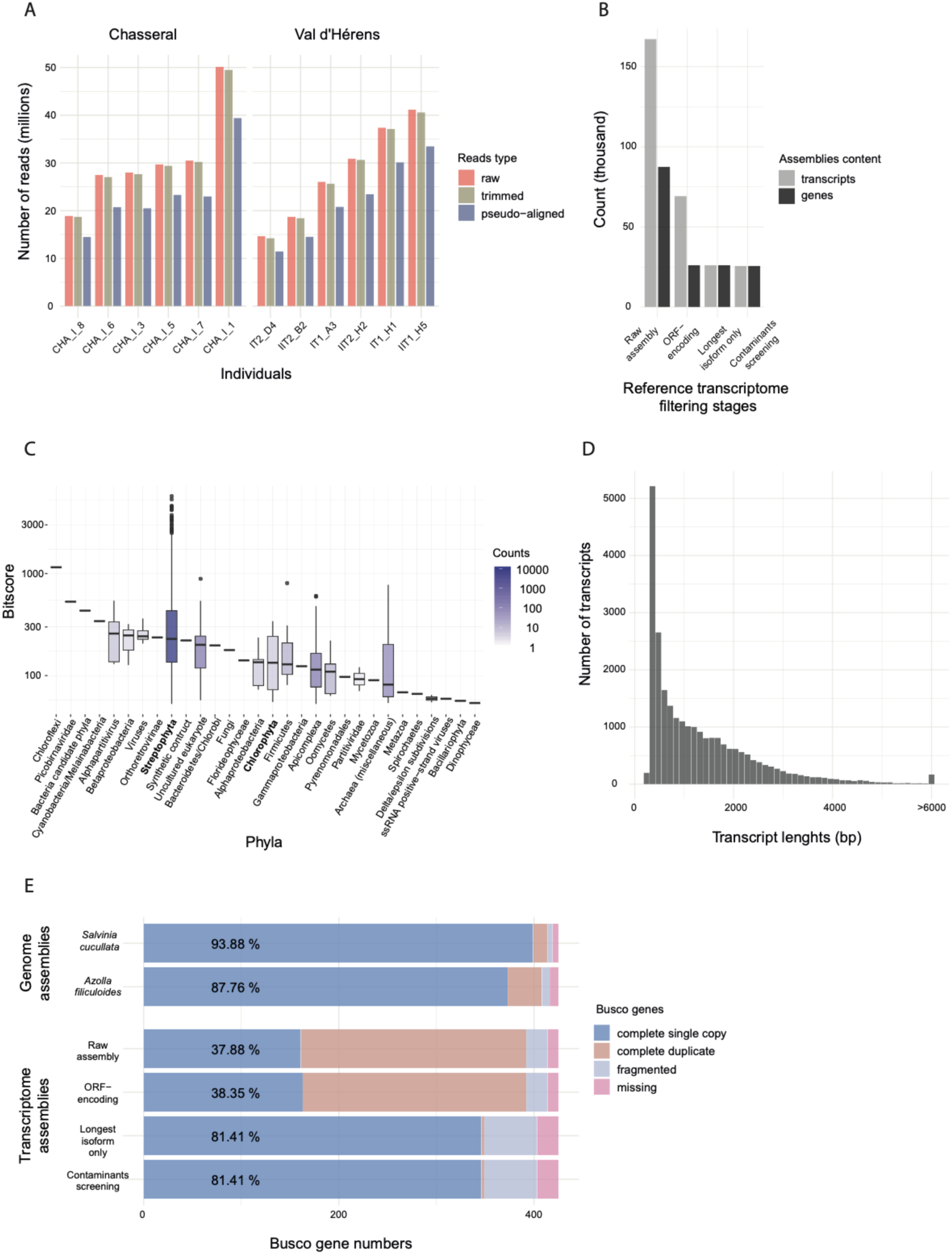
De novo assembly of the *Botrychium lunaria* transcriptome. (A) Distribution of the raw, trimmed, and pseudo-aligned read numbers per individual. (B) Transcripts and gene assembly content at each filtering stage of the transcriptome. (C) Contaminant transcript sequences detected by phylum. Retained phyla are indicated in bold. (D) Distribution of the assembled transcript lengths. (E) BUSCO genes detected at each filtering stage of the *B. lunaria* transcriptome and for two genome assemblies of ferns (*Azolla filliculoides* and *Salvinia cucullata*; Li *et al*., 2018).

**Figure 2:**
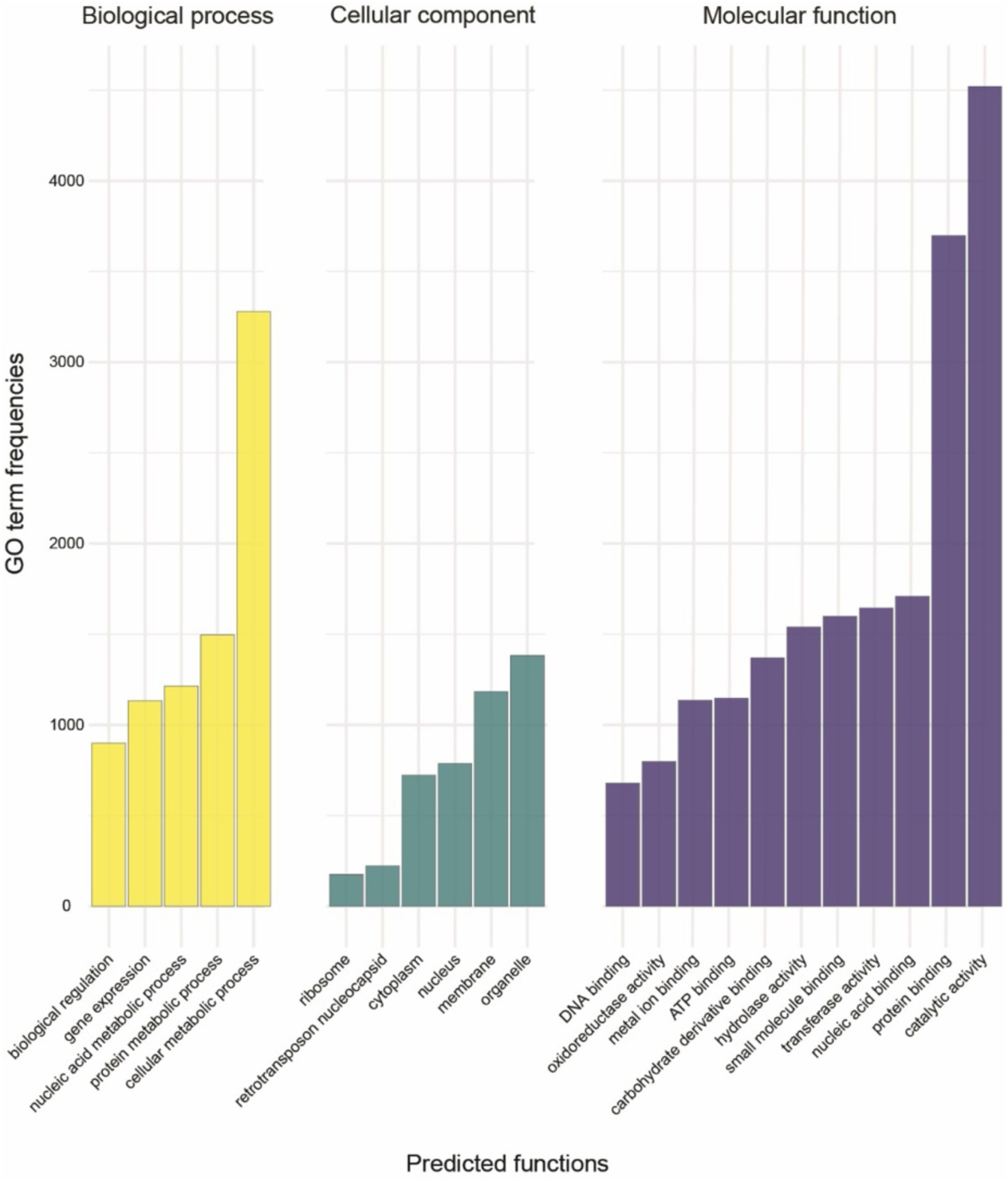
Characterizations of predicted functions encoded by the transcriptome. Gene ontology (GO) term annotations are shown for the 30 most frequent terms per ontology (biological process = BP, cellular component = CC, and molecular function = MF). GO terms with highly similar functions are excluded from the representation.

### Completeness of the B. lunaria transcriptome

We assessed the completeness of the assembled *B. lunaria* transcriptome using BUSCO. Importantly, none of the 30 species constituting the BUSCO viridiplantae_odb10 database belongs to ferns *lato sensu*. We found 81.4% complete single-copy, 0.7% complete duplicates, 12.7% fragmented and 5.2% missing genes for the *B. lunaria* transcriptome (Figure 1E, Table 3). This is comparable to the only two complete genome assemblies of ferns that include *A. filiculoides* with 87.8% and *S. cucullata* with 93.9% complete single-copy genes (Figure 1E, Table 3). The two Salviniaceae genomes and the *B. lunaria* transcriptome exhibited a comparable number of missing BUSCO genes (2.1%, 1.4% and 5.2%, respectively; Figure 1E, Table 3). The *B. lunaria* transcriptome showed a higher percentage of fragmented BUSCO genes compared to genome assemblies of *A. filiculoides* and *S. cucullata* (12.7%, 1.9% and 1.2%, respectively; Figure 1E, Table 3). The mapped reads coverage depth of the reference individual to the assembled transcripts is on average 1,649X with a range of 4 to 514,622X. Most of the transcripts exhibited a moderate coverage (Figure 3A) whereas few (6.9%) showed a coverage > 4,004. The read coverage shows no clear association with transcript length (Figure 3B).

**Table 3:**
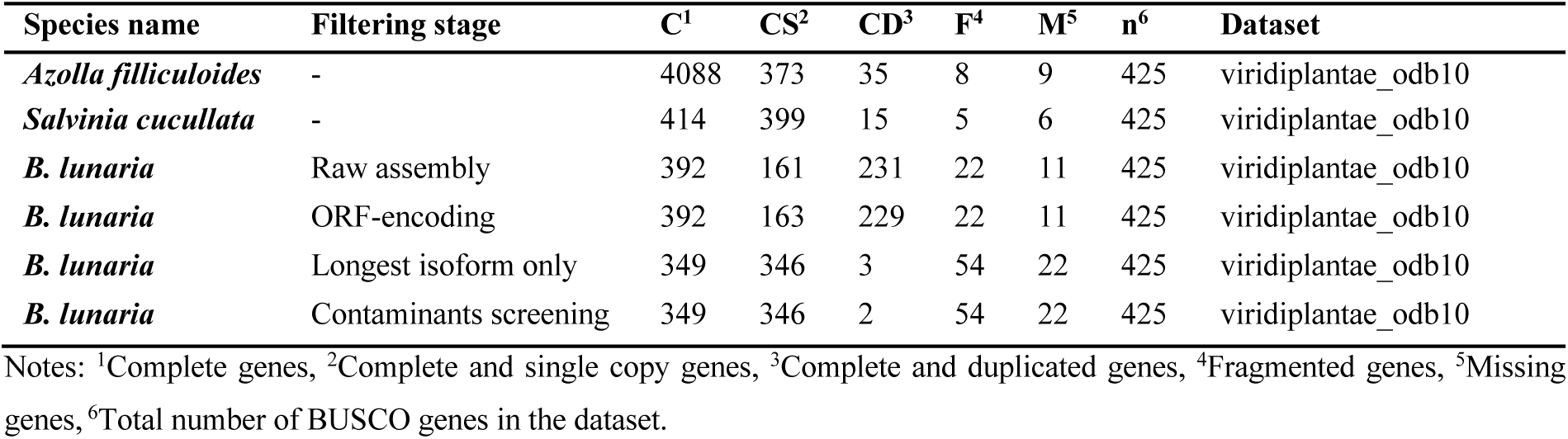
Analyses of assembly completeness using BUSCO genes. The *B. lunaria* transcriptome is compared to two genome assemblies of ferns (*Azolla filliculoides* and *Salvinia cucullata*).

**Figure 3:**
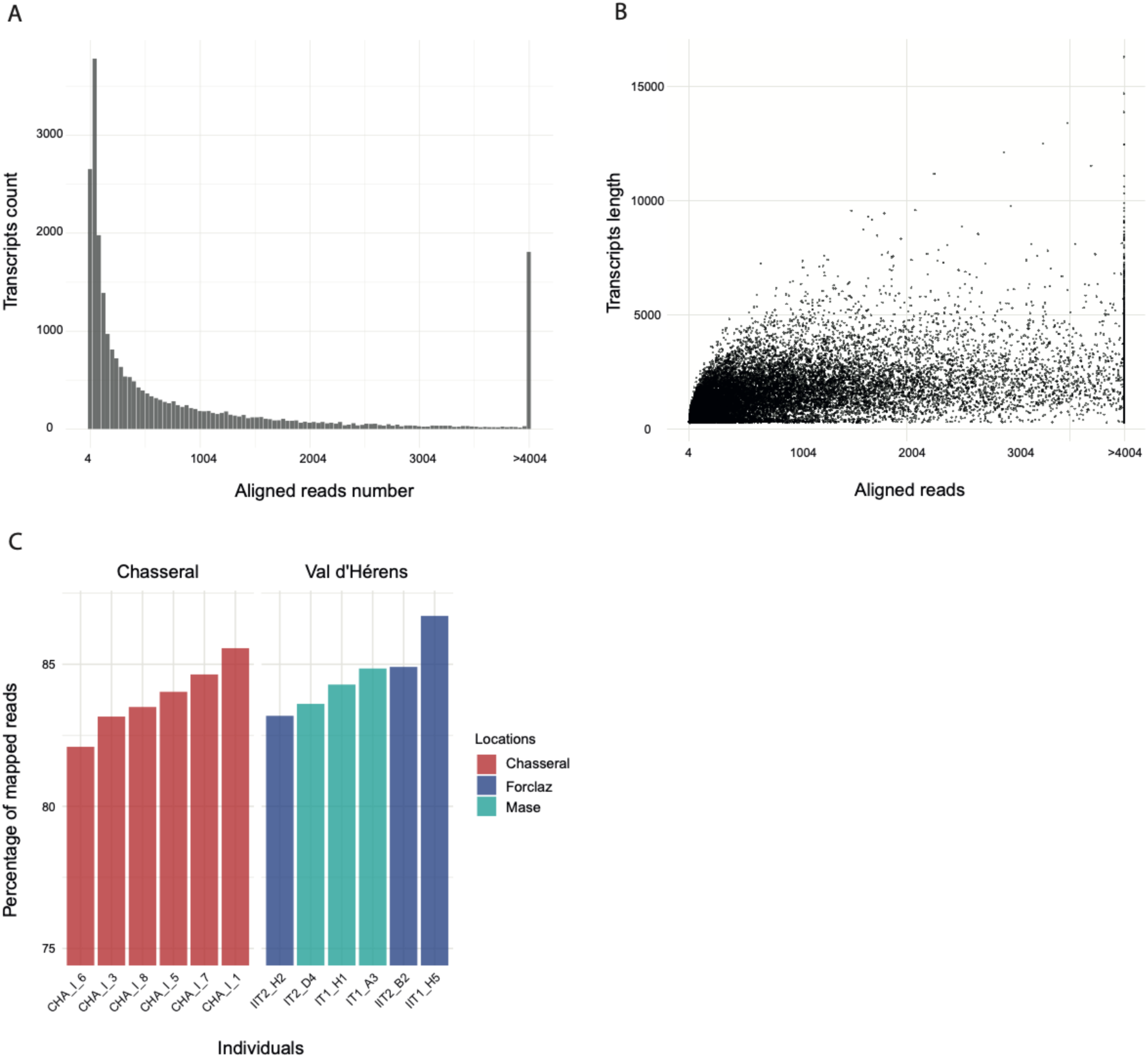
Analyses of the assembly coverage. (A) Number of aligned reads per assembled transcript for the reference individual. (B) Aligned reads of the reference individual according to the assembled transcript length. (C) Mapping rate of all 12 individuals from the Chasseral and Val d’Hérens populations (including the subpopulations Forclaz and Mase). The reference individual used to assemble the transcriptome was CHA_I_1.

### Identification of within-species transcriptomic polymorphism

We mapped reads from the twelve sequenced transcriptomes to the newly established reference transcriptome to identify segregating variants within the species (Figure 3C). The mapping rate of each individual varied between 82.1-86.7%. We found no meaningful difference in mapping rates among populations and individuals. The highest mapping rate was found for the individual IIT1_H5 (86.7%), which was slightly higher than the mapping rate of the reference individual used to establish the transcriptome (CHA_I_1, 85.5%; Suppl. Table S2). Based on reads aligned against reference transcripts, we called SNPs and genotyped each individual as a diploid. Allele frequency distributions show a clear peak around a frequency of 0.5 without secondary peaks at 0.25 or 0.75 (Figure 4A). Hence, all individuals are most likely diploids as higher levels of ploidy would likely have generated additional, minor peaks. We recovered a total of 376,526 high-quality bi-allelic SNPs after filtering. The average number of SNPs per transcript was 17 and the maximum number was 257 (Figure 4B). The SNP density per transcript had a mean of 14, a median of 10, and a maximum of 153 SNPs per kb (Figure 4C). The median SNP density increased with transcript length (Figure 4D).

**Figure 4:**
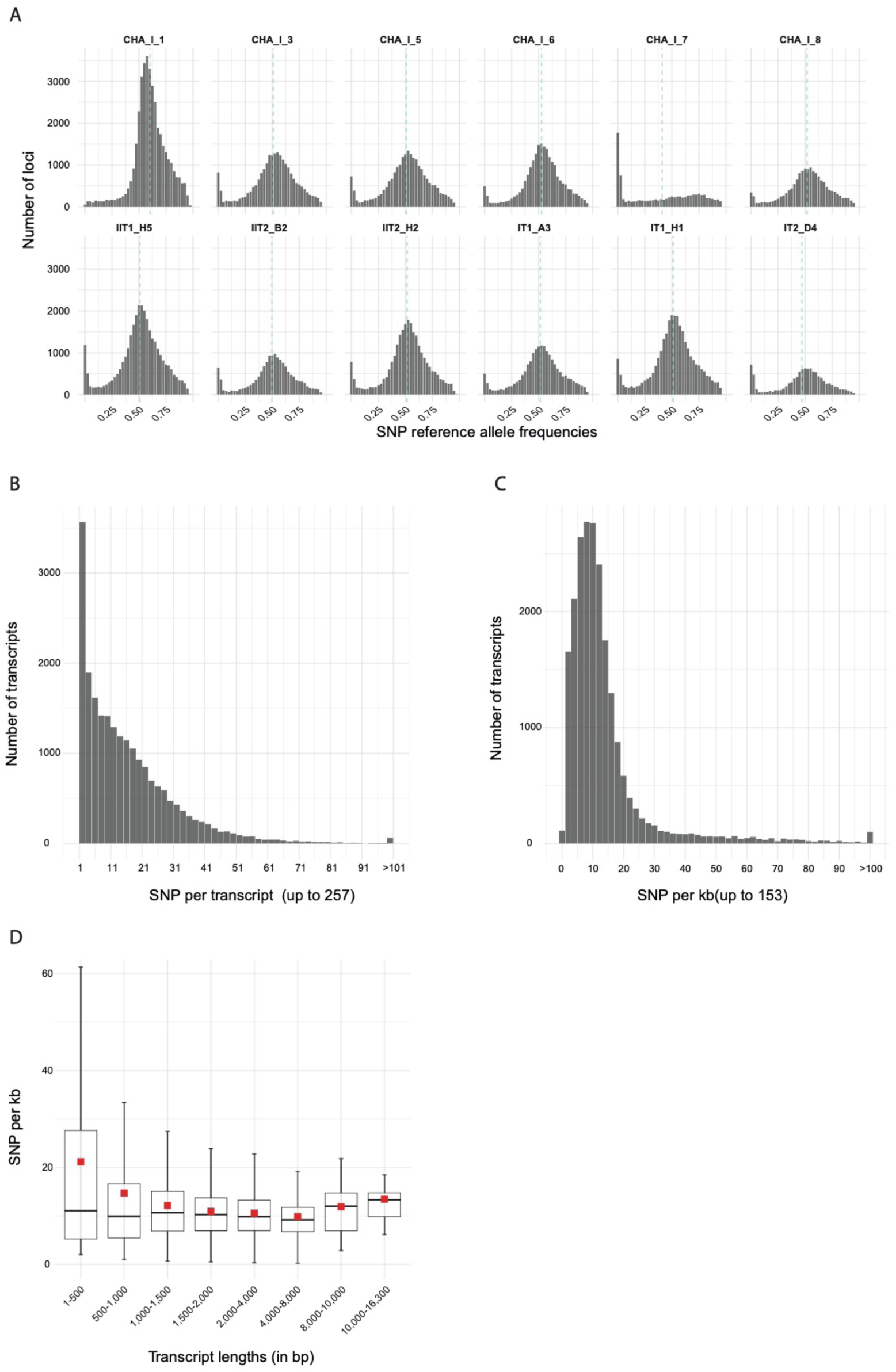
Analyses of population-level transcriptomic polymorphism. (A) Distribution of the transcriptome-wide SNP reference allele frequencies per individual estimated from mapped reads. The light-blue dashed lines show the mean reference allele frequency. Homozygous positions (frequencies 0 and 1) were excluded. (B) Number of SNPs per transcript. (C) Density of SNPs per transcript (i.e., number of SNPs per kb). (D) SNP density according to bins of transcript length. The mean density is shown by a red rectangle.

### Population structure and heterozygosity

We used the SNP genotyping data from the twelve individuals to assess the degree of population differentiation between the sampling locations (Figure 5A). The first principal component (PC1, 17%) of the principal component analysis (PCA) identified a divergent genotype in the Chasseral population (CHA_I_7, Figure 5B). The second principal component (PC2, 12%) separated the two populations Chasseral and Val d’Hérens (Figure 5B). We performed a second PCA excluding the CHA_I_7 individual and found the Chasseral population to be more diverse than Val d’Hérens (Figure 5C). We found no apparent differentiation between the two locations sampled in Val d’Hérens (Forclaz and Mase). The pairwise *F*ST between populations was low (0.079). The mean heterozygosity was slightly higher in Val d’Hérens (*H*e = 0.20) than in the Chasseral population (0.17; Figure 5D). We found similar levels of variation in individual heterozygosity among populations ranging from 0.16 to 0.21, except for CHA_I_7, which was an outlier on the PCA (Figure 5E). CHA_I_7 showed less than half the heterozygosity (*H*e = 0.05) compared to other members of the same population.

**Figure 5:**
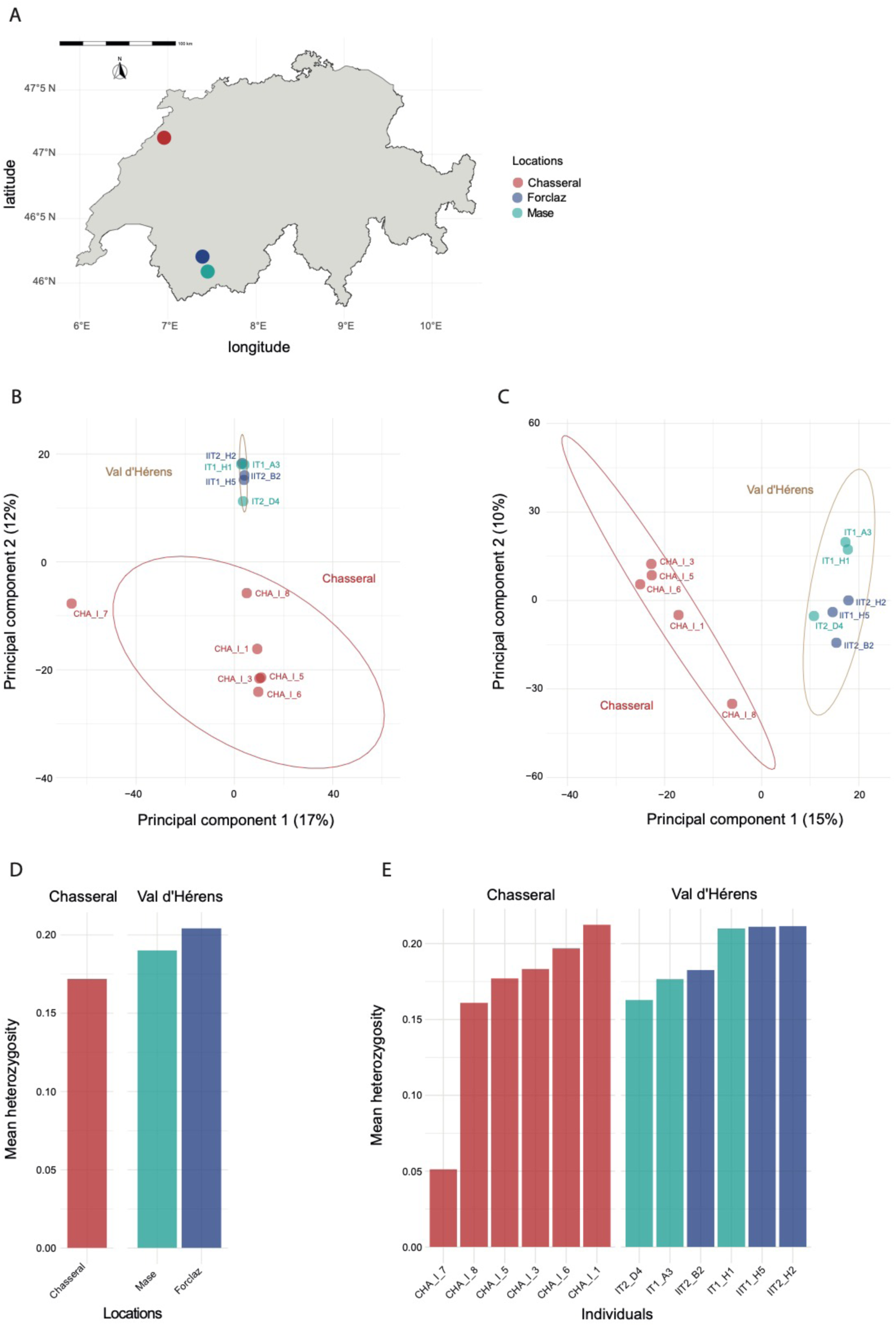
Population genetic structure and observed heterozygosity. (A) Principal component analysis (PCA) of the populations Chasseral and Val d’Hérens (sites Mase and Forclaz). (B) PCA of both populations excluding the CHA_I_7 outlier. Both PCA were analyzed using a reduced SNP dataset of a maximum of 1 SNP per kb of transcript. (C) Mean observed heterozygosity per location grouped by population. (D) Mean observed heterozygosity per individual grouped by population.

### *Phylogenetic inference among* Botrychium *species*

We analyzed available multi-locus sequence data to place the twelve individuals analyzed at the transcriptome level. Among the four nuclear loci previously sequenced in a broad sample of *Botrychium* species, three loci are known to display sequence variation nearly exclusively in intronic sequences (Dauphin *et al*., 2018). Hence, no comparisons with our transcriptomic sequences were possible. We focused on the locus CRY2cA carrying enough informative sites in the coding regions to produce a well-supported phylogeny. The combined dataset for CRY2cA included 67 individuals representing 38 *Botrychium* taxa and an outgroup constituted by *Sceptridium multifidum* and a *Botrypus virginianum* (Dauphin *et al*., 2018; Suppl. Table S3). The multiple sequence alignment contained a total of 3,579 sites and 153 patterns. The main clades Lanceolatum, Lunaria, and Simplex-Campestre were resolved as being monophyletic (Figure 6). Lanceolatum was resolved as a sister group to the Simplex-Campestre and Lunaria clade. All individuals from the Chasseral and Val d’Hérens grouped with *B. lunaria* var. *lunaria* and formed a well-supported clade.

**Figure 6:**
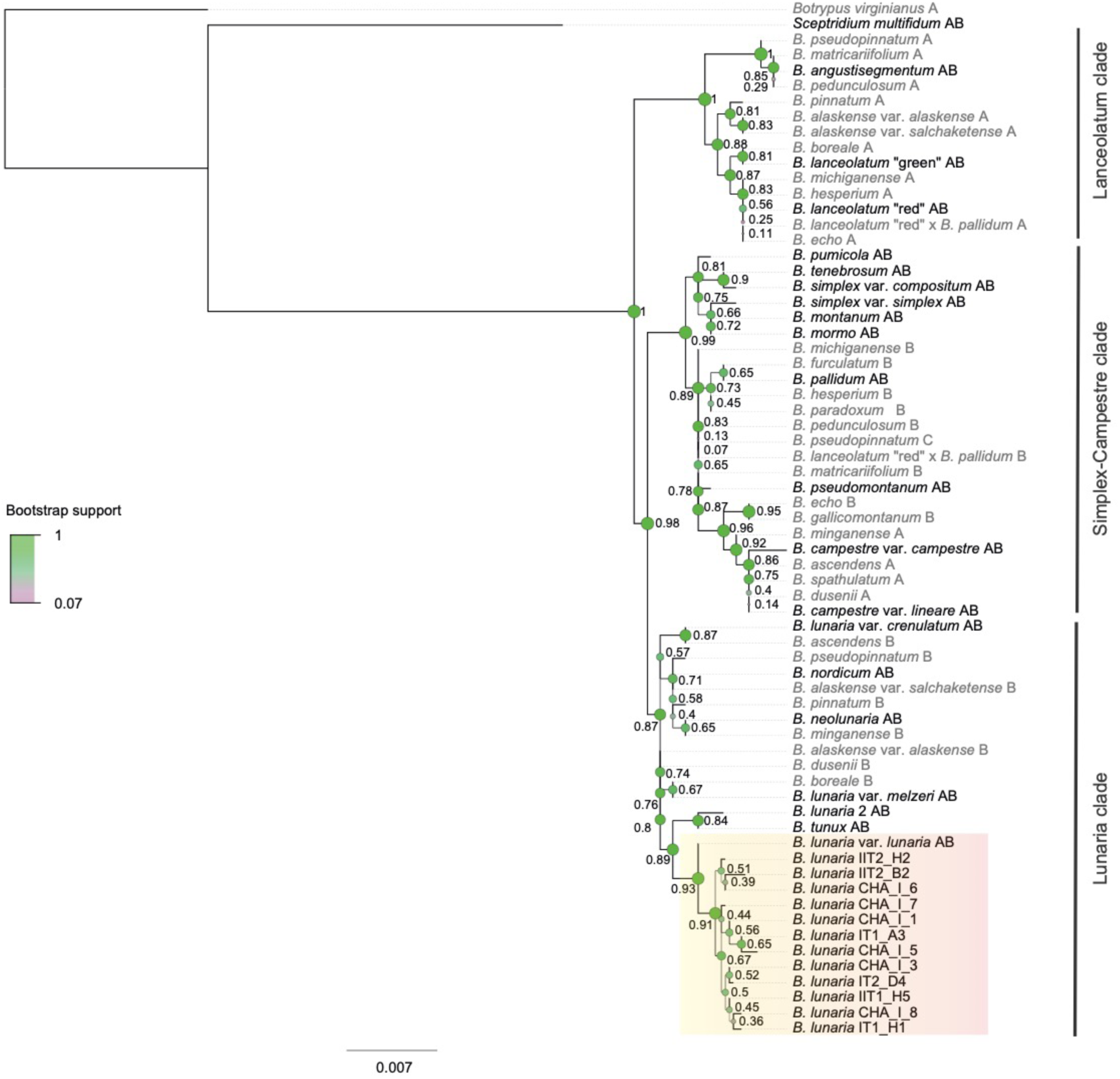
Phylogenetic positions of the *Botrychium lunaria* samples from Chasseral and Val d’Hérens within *Botrychium* genus. Maximum likelihood (ML) tree of the CRY2cA locus. ML bootstrap support values are shown next to nodes. Thicker branch lines, larger node sizes and nodes colored in green further indicate higher bootstrap values. Diploid taxa are shown in black and polyploids are shown in grey. For polyploids, the subgenome is shown individually on the tree and specified by a letter after the species name: “A” for maternal subgenome, “B” for paternal subgenome, and “C” for additional subgenome for the hexaploid (modified from Dauphin *et al*., 2018). The three *Botrychium* main clades were delimited by vertical dark grey lines. Individuals from Chasseral and Val d’Hérens populations are highlighted by an orange rectangle.

### Transcriptome-wide phylogenomic tree of ferns

We retrieved complete transcriptomes of 93 species covering the phylogenetic breadth of ferns (Shen *et al*., 2018; Qi *et al*., 2018; Leebens-Mack *et al*., 2019) in order to robustly place the *B. lunaria* transcriptome. We identified 41,017 orthogroups in total of which we retained 525 orthogroups and 90 species to construct a phylogenomic tree. The species tree branch support values from the local posterior probability (LPP) of the main topology and bootstrap (BS) were highly congruent (Figure 7, Suppl. Figure S2). Minor discrepancies were found in the relationship between two Marattiidae species and the deep relationships among eupolypods. The species tree topology was consistent with the most recent fern phylogenies (Rai and Graham, 2010; Kuo *et al*., 2011; Rothfels *et al*., 2015; Lu *et al*., 2015; Knie *et al*., 2015; Testo and Sundue, 2016; Shen *et al*., 2018; Qi *et al*., 2018; Leebens-Mack *et al*., 2019) and with the current consensus classification (PPG I, 2016). Among the earliest divergent ferns (i.e., eusporangiate and early leptosporangiate), we identified Equisetales as the sister clade to all other ferns and Marattiales as the sister clade to all leptosporangiates (both with 100% LLP and BS support). Hymenophyllales and Gleicheniales were recovered as a monophyletic clade with Dipteridaceae as the sister clade. The Dipteridaceae position was only moderately supported (i.e., LLP = 75, BS = 74). Within the Polypodiales, both the placement of Dennstaedtiineae as sister to all eupolypods and Aspleniaceae as sister to all eupolypods II were strongly supported (i.e., LLP = 1, BS = 1). The quartet support across the tree highlighted poorly resolved branching discussed above (Suppl. Figure 3). Deeper eupolypds relationships remained largely unresolved in our phylogeny. The *B. lunaria* transcriptome clustered with the sister genus *Sceptridium* and the closely related genus *Botrypus*.

**Figure 7.**
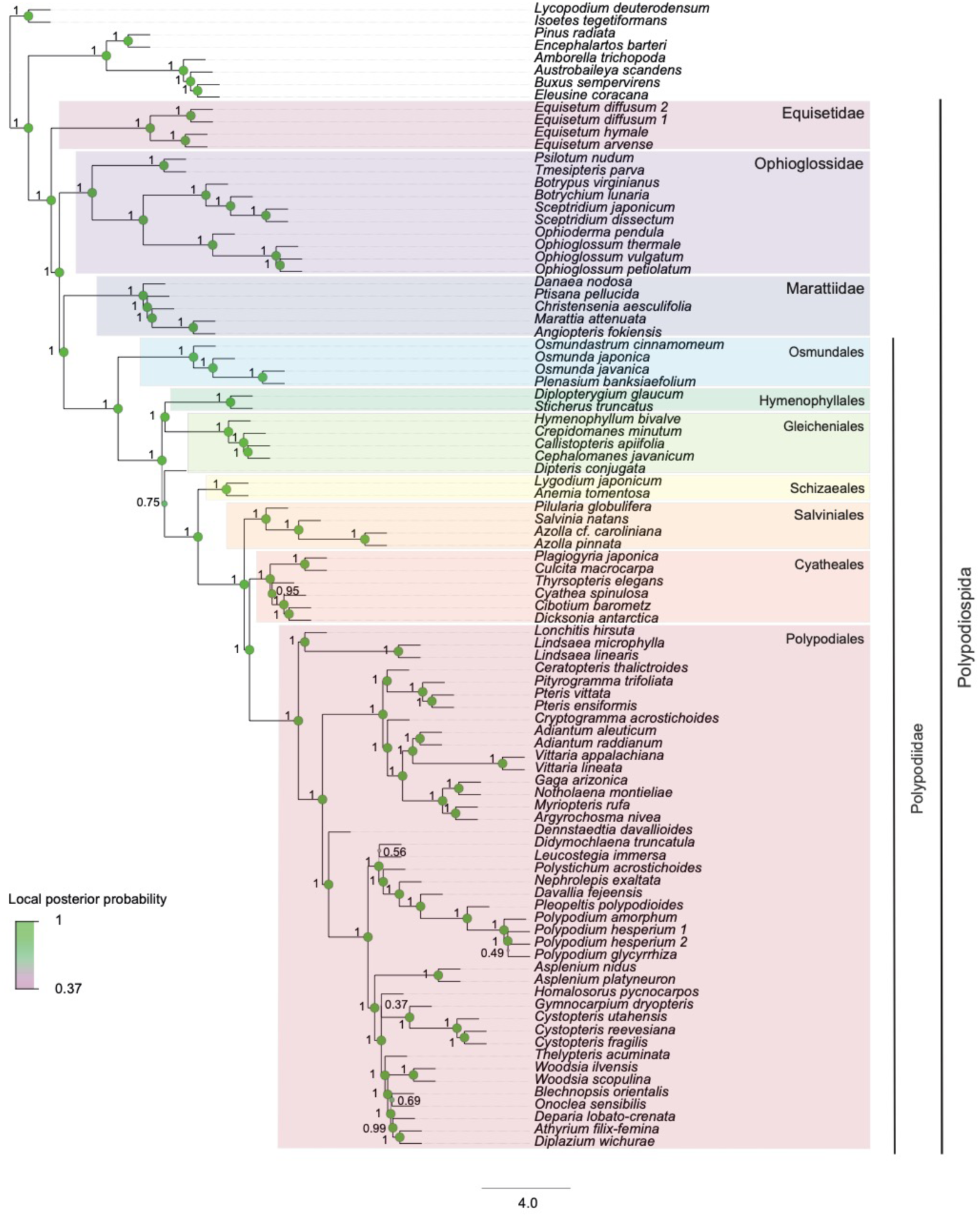
Phylogenomic relationships among ferns including *Botrychium lunaria*. A species tree of 525 orthologous genes including 90 taxa inferred by coalescence-based method implemented in ASTRAL. The branch support is indicated by local posterior probability values of the main topology on node sides. Thicker branch lines, larger node sizes and nodes colored in green further indicate higher local posterior probability values. Fern subclasses (Equisetidae, Ophioglossidae, Marattiidae and Polypodiidae) are denoted by colored rectangles or a vertical dark grey line on clade sides. Polypodiidae orders (Osmundales, Hymenophyllales, Gleicheniales, Schizaeales Salviniales, Cyatheales and Polypodiales) are designated by colored rectangles.

## Discussion

We established a high-quality transcriptome for the genus *Botrychium* filling an important gap in the coverage of early-branching ferns. The completeness of the transcriptomic gene space was comparable to well-assembled fern genomes. Using twelve individuals of the same species sampled in two locations, we were able to generate the first dense SNP dataset for *B. lunaria* and early-branching ferns in general. We were also able to anchor the sequenced individuals in the phylogeny of other *Botrychium* species using a nuclear locus. A phylogenomic tree based on 525 orthologous genes confirmed the phylogenetic position of the genus among other ferns.

### Establishment of a transcriptome for the Botrychium genus

Generating a representative transcriptome assembly is challenging because not all genes are expressed in all tissues and life cycle stages. Across the life cycle of ferns gene expression patterns are largely overlapping (Sigel *et al*., 2018), but the covered gene space is usually increased by including multiple target tissues. For *Botrychium*, only the trophophore and the sporophore were adequate tissues for the extraction of RNA since underground tissues are colonized by arbuscular mycorrhizal fungi (AMF) leading to numerous contaminants. Because we did not include sporophore tissue, the assembled transcriptome potentially underrepresents sporogenesis-specific genes. Despite these challenges, our *B. lunaria* transcriptome has a fairly complete gene space in comparison to a wide range of assembled transcriptomes (Der *et al*., 2011; Shen *et al*., 2018; Qi *et al*., 2018; Leebens-Mack *et al*., 2019). It is important to note that database-dependent tools such as BUSCO consistently underestimate transcriptome completeness if the database was compiled without closely related species. The challenge in using BUSCO is exemplified by the absence of Polypodiopsida species in the viridiplantae dataset. The gene space of assembled fern genomes tends to show less fragmented BUSCO genes compared to the *B. lunaria* transcriptome (Li *et al*., 2018). However, the *B. lunaria* transcriptome is consistent with other high-quality fern transcriptomes (Shen *et al*., 2018; Qi *et al*., 2018; Leebens-Mack *et al*., 2019). Missing gene segments in assembled transcriptomes are often caused by uneven read depth among genes or alternative splicing complicating gene recovery. The completeness of the *B. lunaria* transcriptome compared to other fern genomes and transcriptomes provides a powerful tool for phylogenetic and population analyses.

### Fine-grained resolution of population structure

The transcriptome-wide SNPs revealed clear population structure between two *B. lunaria* populations sampled from Switzerland. The differentiation was apparent even when subsampling SNPs contains a maximum of 1 SNP per kb to avoid biases by highly polymorphic transcripts. It was generally assumed *Botrychium* species show no meaningful genetic differentiation within populations (Farrar, 1998; Hauk and Haufler, 1999) or low genetic differentiation among populations (Camacho and Liston, 2001; Swartz and Brunsfeld, 2002; Birkeland *et al*., 2017). However, the absence of genetic differentiation reported by previous studies may well stem from low marker resolution. The transcriptome-wide markers showed every individual was clearly distinct, and populations showed marked differentiation. The Chasseral and Val d’Hérens populations were collected in the Jura Mountains and Valais Alps, respectively. The two sites are 120 km apart and separated by habitats unsuitable for *B. lunaria*. Hence, reduced gene flow and genetic differentiation among populations is expected. We found no indication of genetic substructure among the two locations Mase and Forclaz within the Val d’Hérens valley. This suggests sufficient gene flow at the local scale or recent recolonization at the upper front of the valley, which is consistent with restriction-site associated DNA sequencing-based analyses of the same field sites (Dauphin, 2017). We found no evidence for higher levels of ploidy based on mapped read depths per individual. Consistent with our findings, a recent study of Swiss populations based on allozyme markers found no evidence for fixed heterozygosity (Dauphin *et al*., 2020). However, we cannot exclude the possibility of very recent polyploidization or autopolyploidization events. Leebens-Mack *et al*. (2019) identified putative ancient whole-genome duplication events in the Ophioglassaceae. No evidence was found for more recent duplication events in the *Botrychium* genus (Dauphin *et al*., 2018).

Population-level genetic diversity is indicative of the reproductive mode of *Botrychium* species populations. Self-fertilization is common and includes sporophytic and gametophytic selfing. In sporophytic selfing, zygotes are produced by gametes from two distinct gametophytes but originate from a single sporophyte. In contrast, in gametophytic selfing, zygotes are produced from gametes of the same gametophyte. Gametophytic selfing is thought to be the main reproductive mode for the *Botrychium* genus and can lead to completely homozygous plants in one generation (Haufler *et al*., 2016). In a population undergoing largely gametophytic selfing, very low genetic variation would be expected among individuals. Hence, the unique genotypes found in the Chasseral and Val d’Hérens populations suggest populations undergo either sporophytic selfing or outcrossing. Interestingly, one of the six Chasseral individuals exhibited less than half the heterozygosity observed in other individuals indicative of recent gametophytic selfing. The genetic diversity of *B. lunaria* populations and the clear structure among sites suggest that sporophytic selfing or outcrossing was dominant with a likely recent gametophytic selfing event. These findings contrast with the general assumption that gametophytic selfing is the dominant reproductive mode in the genus.

### A refined phylogenetic placement of the Botrychium genus

The *B. lunaria* transcriptome enables strong phylogenetic inference at different taxonomic levels overcoming challenges associated with the small number of nuclear and chloroplast markers. *Botrychium* taxa cannot be easily delineated by morphological characteristics, hence taxonomy relies largely on phylogenetics (Dauphin *et al*., 2014; Maccagni *et al*., 2017; Dauphin *et al*., 2017). We have placed the individual *B. lunaria* transcriptomes among other closely related taxa by retrieving orthologous genes, which were previously used for phylogenetic analyses. Despite challenges of low polymorphism in coding sequences of the loci, we were able to recapitulate the phylogenetic position of the reference individual used for transcriptome assembly and the 11 other *B. lunaria*. The newly established transcriptome will enable powerful genome-wide studies across the *Botrychium* genus. Importantly, markers developed using the transcriptome assembly will help to retrace the evolution of the extensive ploidy variation among *Botrychium*.

As an expansion of the phylogenetic analyses within *Botrychium*, we analyzed orthologous genes across all ferns. The genus *Botrychium* was placed within the Ophioglossales with strong support. Furthermore, the phylogenomic tree support the placement of the Marattiidae as a sister clade of the Polypodiidae. The placement of the Marattiidae has long been debated though (Pryer *et al*., 2001; K. M. Pryer *et al*., 2004; Schuettpelz *et al*., 2006; Schuettpelz and Pryer, 2007; Qiu *et al*., 2007; Rai and Graham, 2010; Grewe *et al*., 2013; Wickett *et al*., 2014; Rothfels *et al*., 2015; Lu *et al*., 2015; Knie *et al*., 2015; Testo and Sundue, 2016; Shen *et al*., 2018; Qi *et al*., 2018; Leebens-Mack *et al*., 2019). The uncertain position of the Marattiidae could stem from variation in taxon sampling of the eupsorangiate ferns (Rothfels *et al*., 2015). Phylogenies incorporating broader taxon samples recovered Marattidae as sister to all leptosporangiates with strong support (Rai and Graham, 2010; Knie *et al*., 2015; Rothfels *et al*., 2015; Lu *et al*., 2015; Testo and Sundue, 2016; Qi *et al*., 2018): However, quartet scores in our analyses highlight the remaining uncertainties reported by Leebens-Mack *et al*. (2019) (Suppl. Figure 3). The potential paraphyly observed for the Gleicheniales in our species tree corroborate recent findings based on phylotranscriptomics (Shen *et al*., 2018; Qi *et al*., 2018). Sparse sampling can strongly influence tree topologies. For example, Matoniaceae, constituting one of the tree Gleicheniales families (PPG I, 2016), are not represented in phylogenomics studies. Previous phylogenies based on few barcoding loci only, but with a more representative sampling, identified Gleicheniales as being monophyletic (Pryer *et al*., 2004; Schuettpelz *et al*., 2006; Schuettpelz and Pryer, 2007). The high confidence around the monophyly between the remaining Gleicheniales and the Hymenophyllales suggests a scenario of paraphyly (Figure 7, Suppl. Figure 3). However, phylogenomics studies including Matoniaceae will be need to ascertain the placement of the Dipteridaceae. The identification of Aspleniaceae as the crown group of eupolypods II matches recent studies (Testo and Sundue, 2016; Shen *et al*., 2018) but contrasts with multiple concurrent studies identifying Cystopteridaceae as the crown group (Schuettpelz and Pryer, 2007; Kuo *et al*., 2011; Rothfels *et al*., 2012; Qi *et al*., 2018). Eupolypods II families are notorious for exhibiting family-level heterogeneity in rates of molecular evolution (Rothfels *et al*., 2012; Testo and Sundue, 2016). In our species tree, Aspleniaceae showed long branches compared to other closely related taxa indicative of an accelerated evolutionary rate. Our placement of *Asplenium* may well be caused by the scarce representation of eupolypods II taxa in our dataset and corresponding long branch attraction effect.

Our study establishes a high-quality transcriptome for the early diverging fern genus *Botrychium*. With a genome size estimated at 1C value of 12.10 pg for *B. lunaria* (Veselý *et al*., 2012), the assembly of transcriptomes provides the only currently feasible approach to generate extensive genome-wide markers information. Our phylotranscriptomics analyses identify the *Botrychium* genus as one of the early diverging ferns matching previous phylogenetics analyses on barcoding markers. Furthermore, the inclusion of the *Botrychium* transcriptome improves the resolution of basal nodes among ferns. The transcriptomic markers will be powerful tools to investigate mating systems and polyploidization events. Furthermore, the transcriptome enables fine-grained demographic history analyses helping to dissect evidence for local adaptation across the diverse habitats of ferns.

## Acknowledgements

We thank Frederic Sandoz for assistance in the field work. We thank Aria Minder and Sylvia Kobel for for input on laboratory methods. We acknowledge the 1KP sequencing consortium for advance access to transcriptomic sequences. Emilie Chanclud, Ursula Oggenfuss and Erik Koenen provided advice on data analysis. Ursula Oggenfuss, Leen N. Abraham, Simone Fouché, Pierre-Emmanuel Du Pasquier, Rosangela Ston, Erik Koenen and Giacomo Zilio provided helpful comments on a previous manuscript version. Rosangela Ston for mounting the herbarium vouchers. Data produced and analysed in this paper were generated in collaboration with the Genetic Diversity Centre (GDC), ETH Zurich and the Functional Genomics Center (FGC), Zurich. This work was supported by the Overhead fund of the University of Neuchâtel.

## Data availability

Raw sequencing reads and the assembled transcriptome are available at the NCBI Short Read Archive under the Bioproject accession PRJNA605155. Sequences for multi-gene phylogenies are available under the NCBI Nucleotide accession (*pending*). Supplementary files (phylogenetic trees, alignments and protein sequences) are available: https://doi.org/10.5281/zenodo.3959727

