## Supplementary Figures for "A transcriptome for the early-branching fern *Botrychium lunaria* enables fine-grained resolution of population structure"

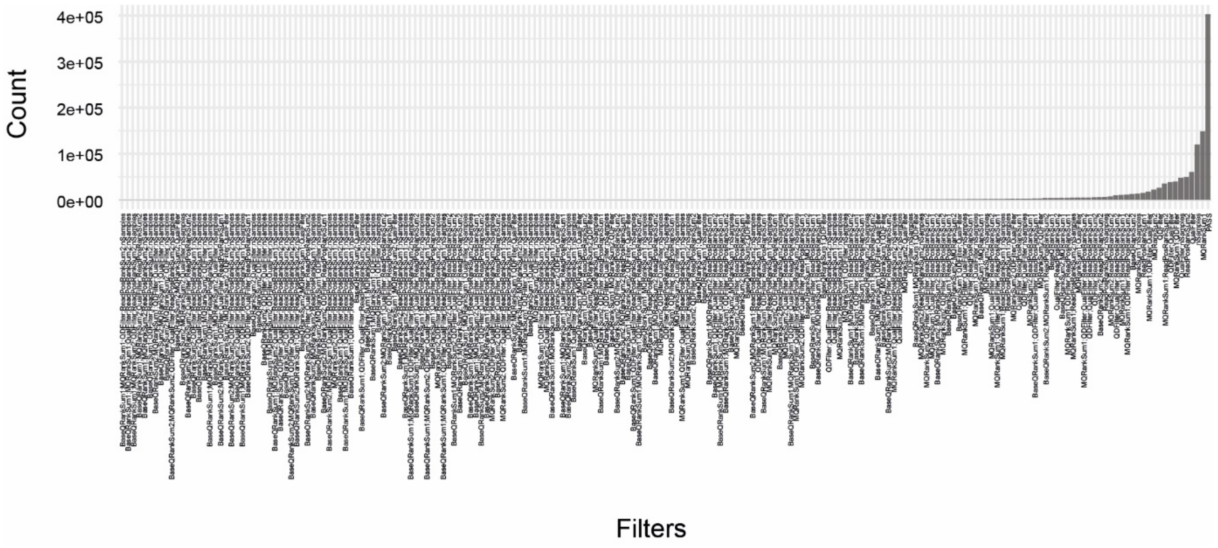


**Supplementary Figure S1: SNP filtering outcome.** The number of filter flagged loci per individual (or combination of quality criteria). PASS identifies the number of SNPs passing all filtering criteria.

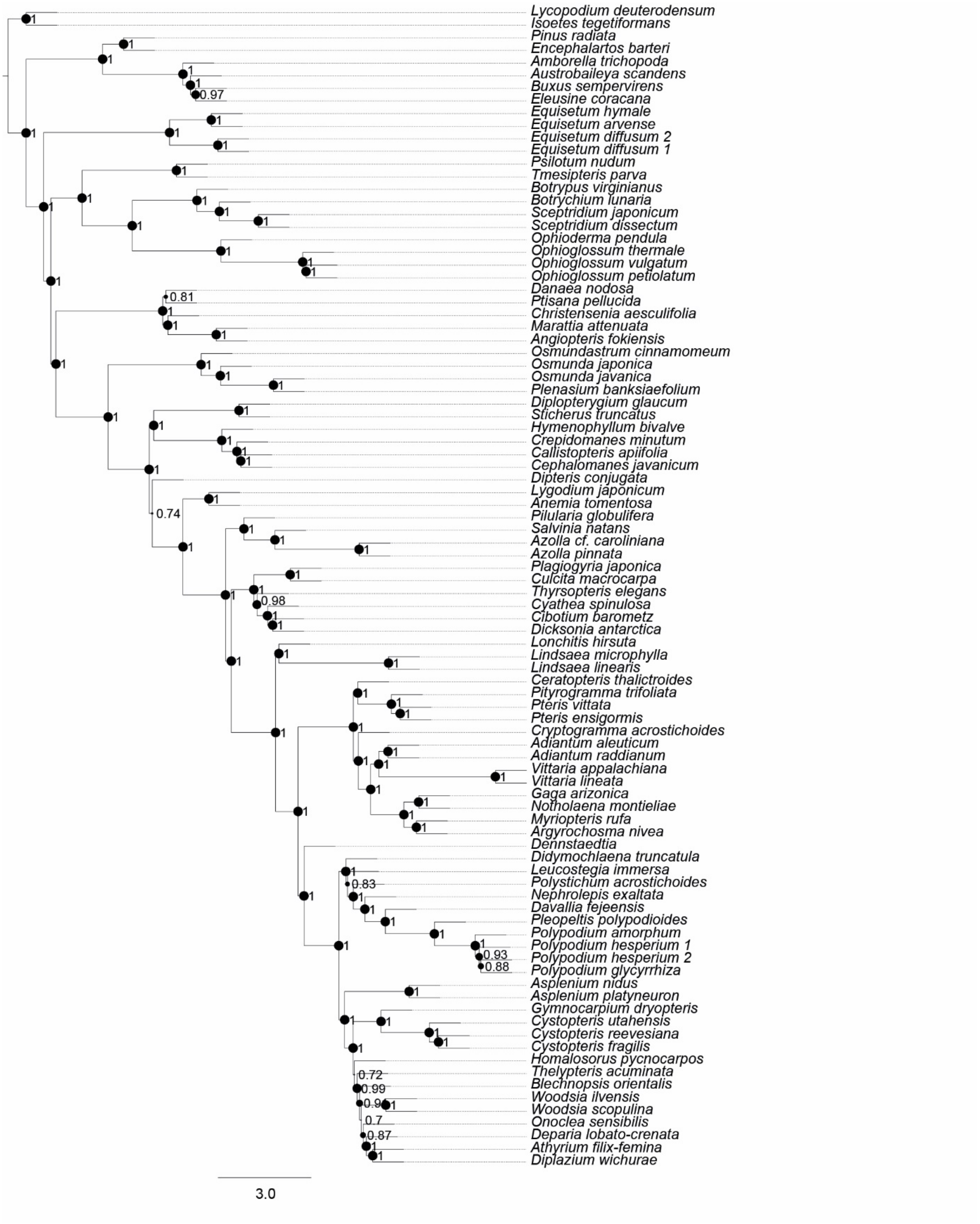


**Supplementary Figure S2: Species tree estimated out of 525 orthologous genes including 90 taxa using Astral.** The branch support is indicated by the bootstrap values at the nodes. Larger node sizes correspond to higher bootstrap values.

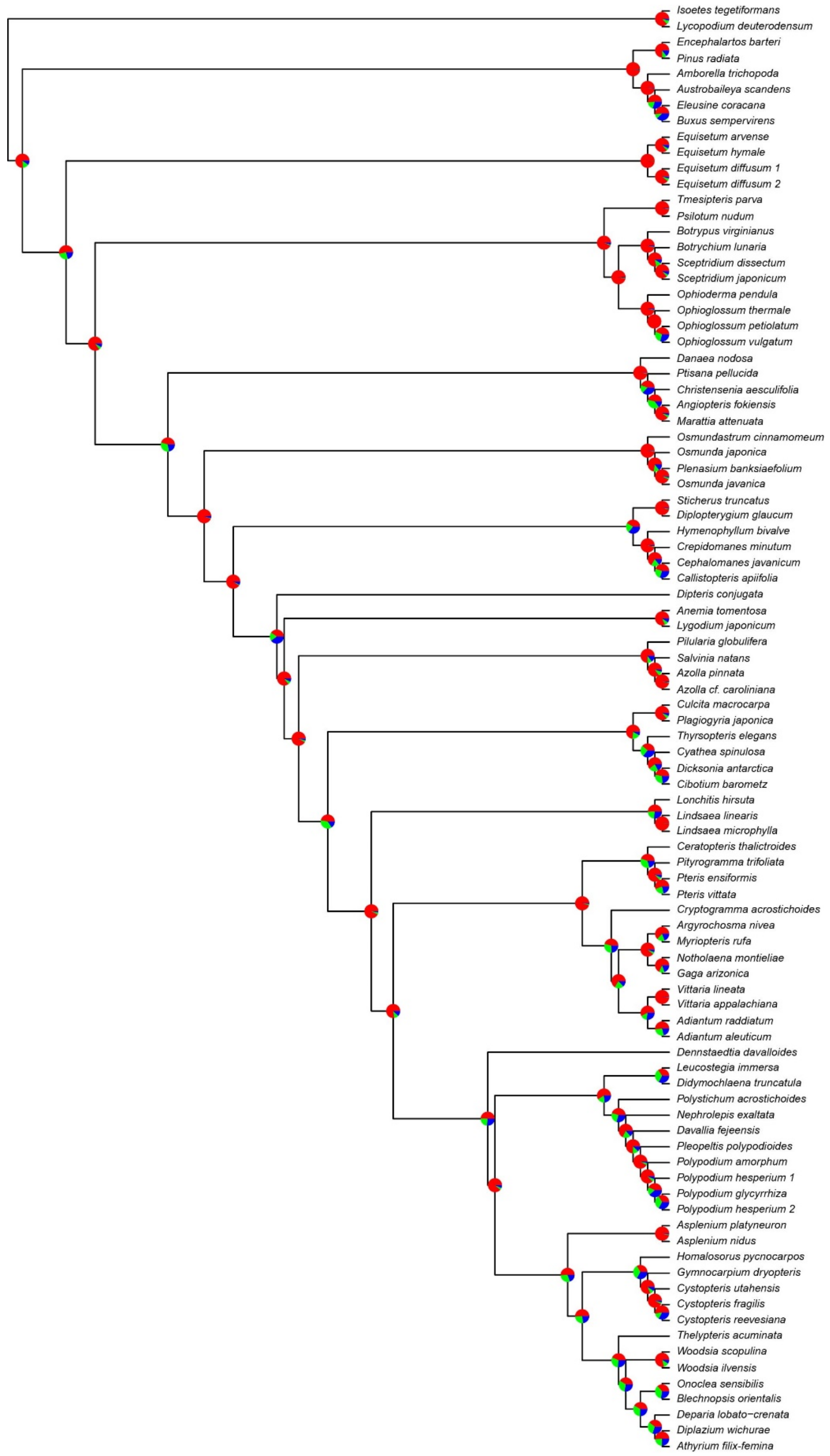


**Supplementary Figure S3: Astral species tree on which the branches quartet support values were depicted by pie charts**. Inside each pie chart, the red, the green and the blue slices correspond to the normalized quartet scores for the main topology, the first and the second alternative, respectively.

### Supplementary Tables

(see separate files)

**Supplementary Table S1: Information about the transcriptomes used in the phylogenomic analyses.** The identifiers refer to those originally published by Leebens-Mack et al. (2019), Qi et al. (2018) and Shen et al. (2017) and used in this study.

**Supplementary Table S2: Raw and trimmed read distribution for the *Botrychium lunaria* transcriptomes.** Percentage of retained reads after trimming, pseudo-aligned reads by Kallisto and reads aligned by Bowtie.

**Supplementary Table S3: Information regarding the CRY2cA sequence dataset used for the phylogenetic inference of Botrychium species.** The individual identifiers refer to those used by Daupin et al. (2018) and in this study.

**Supplementary Table S4**: **Orthogroups table given by OrthoFinder.**

**Supplementary Table S5: Filtered orthogroups table.**

### Supplementary Files

[Available from Zenodo: https://doi.org/10.5281/zenodo.3959727]

**File S1: Alignment of Botrychium CRY2cA sequences used to infer the genus-level phylogeny.**

**File S2: Peptide sequences used to infer the orthogroups named according species names or identifiers** (see Table S1).

**File S3: Alignments of the orthogroup sequences subset used to infer the phylogenomy.**

**File S4: Output files from modeltest-ng named by orthogroup names.**

**File S5: Output files from raxml-ng named by orthogroup names.**
